# RNA polymerase mapping in plants identifies enhancers enriched in causal variants

**DOI:** 10.1101/376640

**Authors:** Roberto Lozano, Gregory T. Booth, Bilan Yonis Omar, Bo Li, Edward S. Buckler, John T. Lis, Jean-Luc Jannink, Dunia Pino del Carpio

**Affiliations:** Plant Breeding and Genetics, School of Integrative Plant Science, Cornell University, Ithaca, NY, USA; Department of Molecular Biology and Genetics, Cornell University, Ithaca, NY, USA; Montpellier SupAgro, 34060 Montpellier Cedex 02, France; State Key Laboratory of Plant Genomics and National Center for Plant Gene Research, Institute of Genetics and Developmental Biology, Chinese Academy of Science, Beijing, China; Institute for Genomic Diversity, Cornell University, Ithaca, NY, USA; United States Department of Agriculture, Agricultural Research Service (USDA-ARS) R.W. Holley Center for Agriculture and Health, Ithaca 14853, NY, USA; Department of Economic Development, Jobs, Transport and Resources, AgriBio Centre for AgriBioscience, Bundoora, Australia

## Abstract

Promoter-proximal pausing and divergent transcription at promoters and enhancers, which are prominent features in animals, have been reported to be absent in plants based on a study of *Arabidopsis thaliana*. Here, our PRO-Seq analysis in cassava (*Manihot esculenta*) identified peaks of transcriptionally-engaged RNA polymerase II (Pol2) at both 5’ and 3’ ends of genes, consistent with paused or slowly-moving Pol2, and divergent transcription at potential intragenic enhancers. A full genome search for bi-directional transcription using an algorithm for enhancer detection developed in mammals (dREG) identified many enhancer candidates. These sites show distinct patterns of methylation and nucleotide variation based on genomic evolutionary rate profiling characteristic of active enhancers. Maize GRO-Seq data showed RNA polymerase occupancy at promoters and enhancers consistent with cassava but not Arabidopsis. Furthermore, putative enhancers in maize identified by dREG significantly overlapped with sites previously identified on the basis of open chromatin, histone marks, and methylation. We show that SNPs within these divergently transcribed intergenic regions predict significantly more variation in fitness and root composition than SNPs in chromosomal segments randomly ascertained from the same intergenic distribution, suggesting a functional importance of these sites on cassava. The findings shed new light on plant transcription regulation and its impact on development and plasticity.

## Main Text

Gene expression in plants is a highly regulated process controlling the production of coding and noncoding RNA molecules and is central to development and phenotypic plasticity. The dynamics of transcriptional regulation have been extensively studied in several model organisms, including humans, yeast, and fruit flies^1^. These studies have revealed a complex network of molecular elements that orchestrate gene expression patterns and thereby shape the transcriptional landscape of each organism. Failure in gene regulation control can have detrimental effects in development and lead to disease^2^.

Nascent RNA sequencing techniques such as Global nuclear Run-On sequencing (GRO-seq)^3^ or Precision nuclear Run-On sequencing (PRO-seq)^4,5^ have been used to map and quantify transcriptionally-engaged polymerase density. These techniques have identified promoter-proximal pausing of Polymerase II (Pol II) and bi-directional transcription^6^ as widespread phenomena in metazoans^7^. The pausing of elongating Pol II occurs shortly after the Pre-Initiation Complex is assembled and initiation has occurred^1^. Promoter-proximal pausing has been suggested as a mechanism to tune the expression of specific genes in response to external regulatory signals and might also play a role in stabilizing the open chromatin state around promoter regions^1^. Nascent RNA sequencing also revealed the presence of bidirectional transcription in enhancers, supporting a more unified model of transcription initiation between enhancers and promoters^8^.

In plants, nascent RNAs have only been profiled in *Arabidopsis thaliana*^*9*^ and *Zea mays*^*10*^ using GRO-seq. In *Arabidopsis*, the analysis revealed a lack of bi-directional transcription, promoter-proximal (i.e., 5’) pausing or expression of enhancer RNAs. Instead, prominent 3’ accumulation of RNA polymerase was observed in both maize and Arabidopsis^9^ These data suggested that gene regulation in plants may have diverged from what is observed in other eukaryotes, and Hetzel et al.^9^ suggested that the presence of Pol IV and Pol V (absent in metazoans)^11^ reflects a different evolutionary approach to gene regulation within the plant kingdom.

We characterized nascent transcription in cassava (*Manihot esculenta*) and maize seedlings using PRO-seq^5,11^ and re-analyzed the *Arabidopsis* and maize GRO-seq data from Hetzel et al.^9^ and Erhard et al.^10^ respectively. For clarity, the four libraries will be referred to as PRO-cassava, PRO-maize, GRO-maize and GRO-arabidopsis. In agreement with previous studies, GRO-arabidopsis lacked promoter proximal pausing (Fig 1A) and, instead, showed accumulation of engaged polymerases at the 3’ end of each gene (Fig 1B). Analysis of PRO-maize showed the previously reported 3’ pausing^9^ (Fig 1C) and a small accumulation of reads at the Transcription Start Site (TSS) (Fig 1B). This accumulation was consistent with GRO-maize, and thus general across the two techniques and two varieties of maize (Fig S1). PRO-cassava, in contrast, showed a clear pattern of both 5’ (Fig 1E) and 3’ pausing (Fig 1F). Out of the 24,532 genes that were expressed in cassava, 16,605 had a Pausing Index (PI) higher than 2 (Fig S2, Supplemental methods). While all three plant species demonstrated polymerase accumulation at the 3’ end of genes, each displayed a unique accumulation pattern in the promoter-proximal region. Unlike mammals, bi-directional transcription was uncommon among plant promoters (Fig 1). However, several cassava genes showing this behavior were identified (Fig S3).

**Fig. 1.**
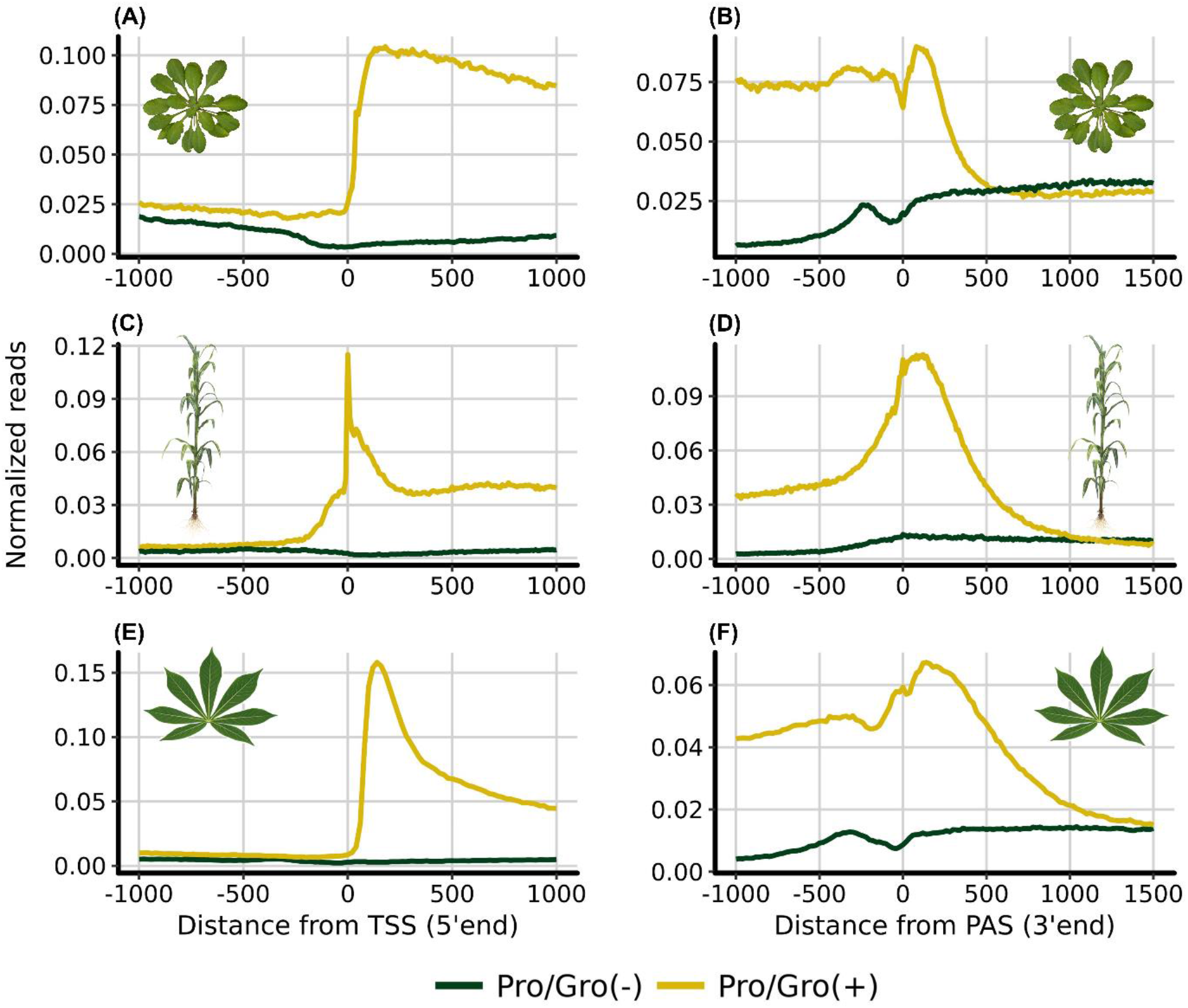
Accumulation of Pro-seq reads around the Transcription Start Site and the Polyadenylation sites of three different plant species. Metaplot of GRO/PRO-seq signal from annotated genes normalized for reads per bp per gene in *Arabidopsis thaliana* (GRO-seq, n = 28,775) (A, B), *Zea mays* (PRO-seq, n = 38,943) (C,D) and *Manihot esculenta* (PRO-seq, n = 31,895). Reads were aligned to the TSS and the PAS in both sense (yellow) and antisense (green) directions relative to the direction of gene transcription (E,F). Prominent promoter-proximal pausing is shown in *Manihot esculenta*, and in some degree in maize, but it is not present at all in Arabidopsis as previously reported^9^. Accumulation of RNA polymerase at the 3’ end of the genes is a common feature in the three plant species.

Enhancers are key eukaryotic regulatory elements that control spatiotemporal gene expression and are especially important during development^12^. Studies in mammals have shown that enhancers produce short unstable RNAs known as eRNAs^13,14^. In the first study of nascent RNA in *Arabidopsis*, aligning GRO-seq reads to open chromatin sites did not identify transcribed plant enhancers^9^ To validate this observation in another species, we mapped PRO-seq peaks outside coding regions (3kb from the 5’UTR or 3’ UTR of any gene) in the cassava genome. We identified ~2,000 peaks in intergenic regions showing clear bi-directional transcription, similar to that observed in mammalian and other metazoan enhancers (Fig S4). Given the resemblance of these elements to mammalian transcriptional regulatory elements, we used discriminative regulatory-element detection (dREG)^15^, a support vector regression algorithm trained to detect enhancers and promoters from GRO-seq mapped reads. We identified 34,000 candidate regulatory regions across the genome of which 16,800 were located in intergenic regions, and 9.665 were at least 1kb away from any gene. This set of 9,665 regions, which we refer to as enhancer candidate regions (Supplemental File S1), showed a clear asymmetric pattern of transcription (Fig 2A-2B).

**Fig. 2.**
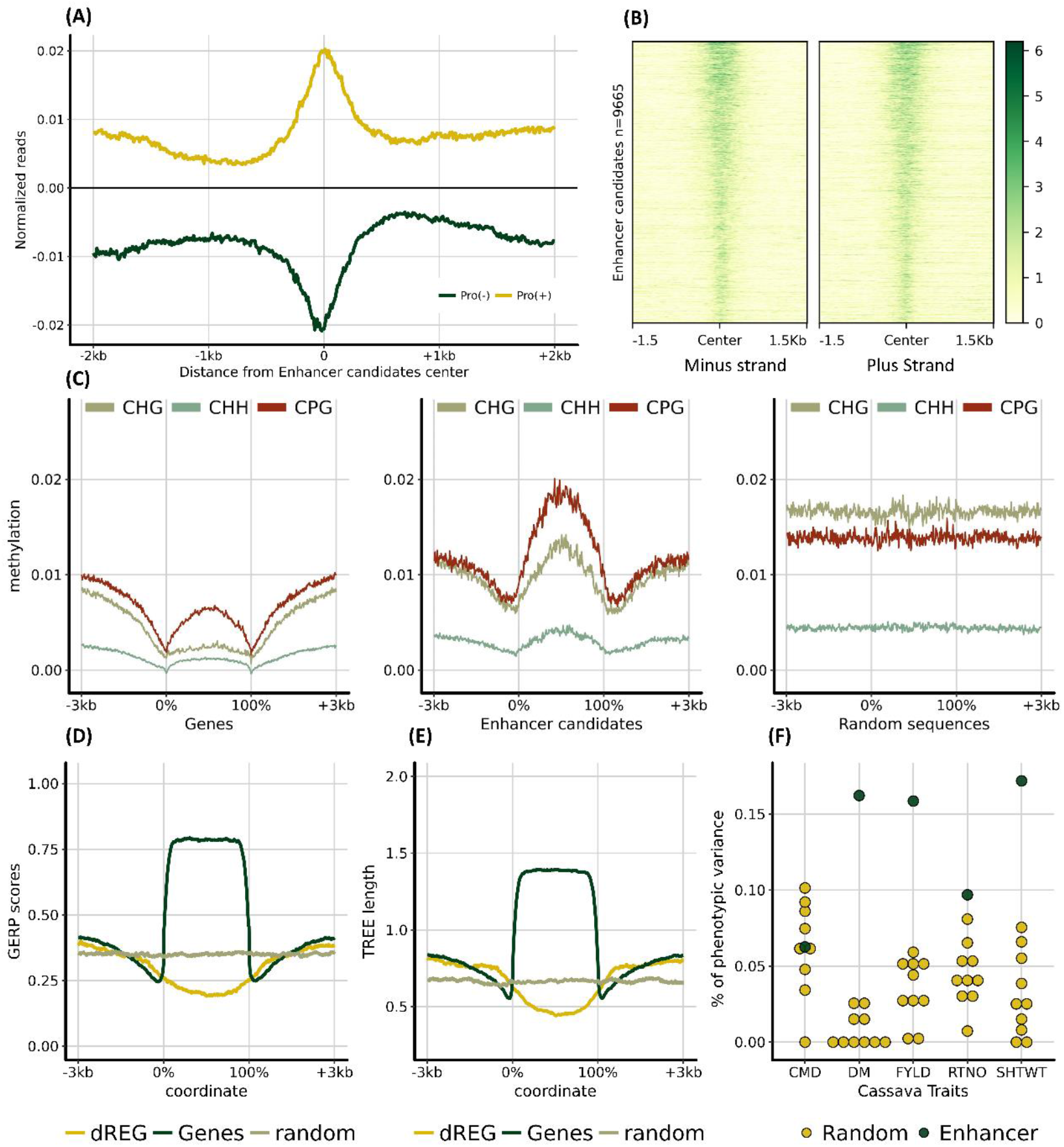
Enhancer candidates in cassava have a particular methylation pattern, are evolutionary less conserved and explain more phenotypic variance than expected for several agronomic traits. (**A**) Pro-seq reads mapping around cassava enhancer candidates. Reads were sorted by strand and the normalize reads were plotted around the center of each candidate. (**B**) Heatmap representation of reads mapping to the enhancer candidate regions. The regions are sorted based on dREG scores. (**C**) Cassava methylation patterns for the three methylation contexts (CpG, CHH and CHG) were plotted around the genic regions, enhancer candidate regions and a set of random sequences. The random set has the same number and length distribution as the enhancer candidates. Genomic regions were scaled (0-100%) for visualization. (**D**) Genomic Evolutionary Rate Profiling (GERP) scores and corresponding tree lengths (**E**) were also plotted around the Enhancer candidates (dREG), Genes and random set of regions. (**F**) Genomic Partitioning of complex agronomic traits (DM: Dry matter content, FYLD: Fresh yield, RTNO: Root number, SHTWT: Shoot weight) and a disease trait (CMD: Severity of Cassava Mosaic Disease). Relationship matrices were calculated using SNP markers within the enhancer candidate regions using the LDAK5 model and variance components were estimated using EMMREML.

Two independent lines of evidence supported the biological activity of these enhancer candidate regions. First, using DNA methylation data previously measured in cassava cultivar TME7^16^, we observed profiles in the three DNA contexts (CpG, CHG, and CHH) around the 9.665 cassava enhancer candidate regions distinct from genic and random regions across the genome (Fig 2C). Second, Genomic Evolutionary Rate Profiling (GERP)^17−19^ of the enhancer candidates showed lower conservation of these regions than of random sequences across the genome, whereas coding regions are conserved (Fig 2D and 2E). This low conservation agrees with observed patterns in mammals^20^ where enhancers, unlike promoters, are rarely conserved and evolve rapidly.

To test the functional relevance of the plant enhancer candidates, we estimated the percentage of the SNP heritability^21^ attributable to the enhancer candidates as compared to randomly ascertained genomic regions with the same intergenic distribution. We set up genomic partitions^22,23^ separating a focal partition from a rest-of-genome (ROG) partition, where the focal partition was either the enhancer candidates or the random regions. We used 3,011 cassava clones (i.e., genetically unique but each clonally propagated) of the NextGen Cassava Breeding Project^24^ evaluated for four quantitative agronomic traits, dry matter content (DM), fresh yield (FYLD), number of roots (RTNO), and shoot weight (SHTWT), and a disease trait, mean cassava mosaic disease severity (MCMDS) whose genetic architecture is controlled primarily by a single resistance locus^25^. Genomic relationship matrices (GRM) for each partition were calculated using LDAK5, following recommendations made by Speed et al.^21^. For each partition, we fit a model estimating variances of effects distributed according to ROG and focal GRMs. The percentage of phenotypic variance explained by markers inside enhancer regions was higher than random regions for all quantitative agronomic traits but not for the disease resistance trait (Fig. 2F). Plant disease resistance is often conditioned by genes that cause recognition of infection rather than by differential expression^26^. In contrast, fitness-related quantitative traits can be strongly affected by gene regulation^27^. Together, these results suggest that the enhancer candidates causally affect plant phenotypes by modulating gene transcription.

In summary, the nascent transcriptome of cassava, as revealed by PRO-seq, showed features of transcriptional regulation that weren’t present or detected in previous plant experiments, including promoter-proximal pausing and the presence of eRNAs. We note that plants, like yeast, lack Negative ELongation Factor (NELF), which is likely required for a kinase-regulated release of paused Pol II^28^. Thus, this enrichment of Pol II may reflect a related maturation checkpoint observed in fission yeast^29^. Enhancer candidate regions were shown to have low levels of evolutionary conservation, a bi-directional pattern of transcription, and a specific DNA methylation profile. These regions also contributed disproportionately to fitness and root composition variation, showing an important new way forward in prioritizing genomic regions for use in crop improvement.

In an attempt to extend our observations to other plants we analyzed the existing GRO-maize data similarly to the PRO-cassava data. We identified 4,135 (Supplemental file S2) enhancer candidate regions using dREG and again observed a clear pattern of bi-directional transcription (Fig 3A, 3B). The transcription of these candidates was conserved across different genotypes as the PRO-maize data generated in our study showed transcription signals in the same regions (Fig. S4). We analyzed the transcription pattern of 1,495 (Supplemental file S3) enhancer candidates identified in a separate maize study^30^ using an approach that integrated genome-wide methylation data, chromatin accessibility (DNAse-seq), and histone marks (H3K9ac). The GRO-maize reads mapped against these candidates also showed bi-directional transcription (Fig 3C,3E). Moreover, 519 (Supplemental file S4) of the 1,495 enhancer candidates (~35%) showed significant levels of bi-directional transcription and were identified by dREG (Fig 3D, 3F).

**Fig. 3.**
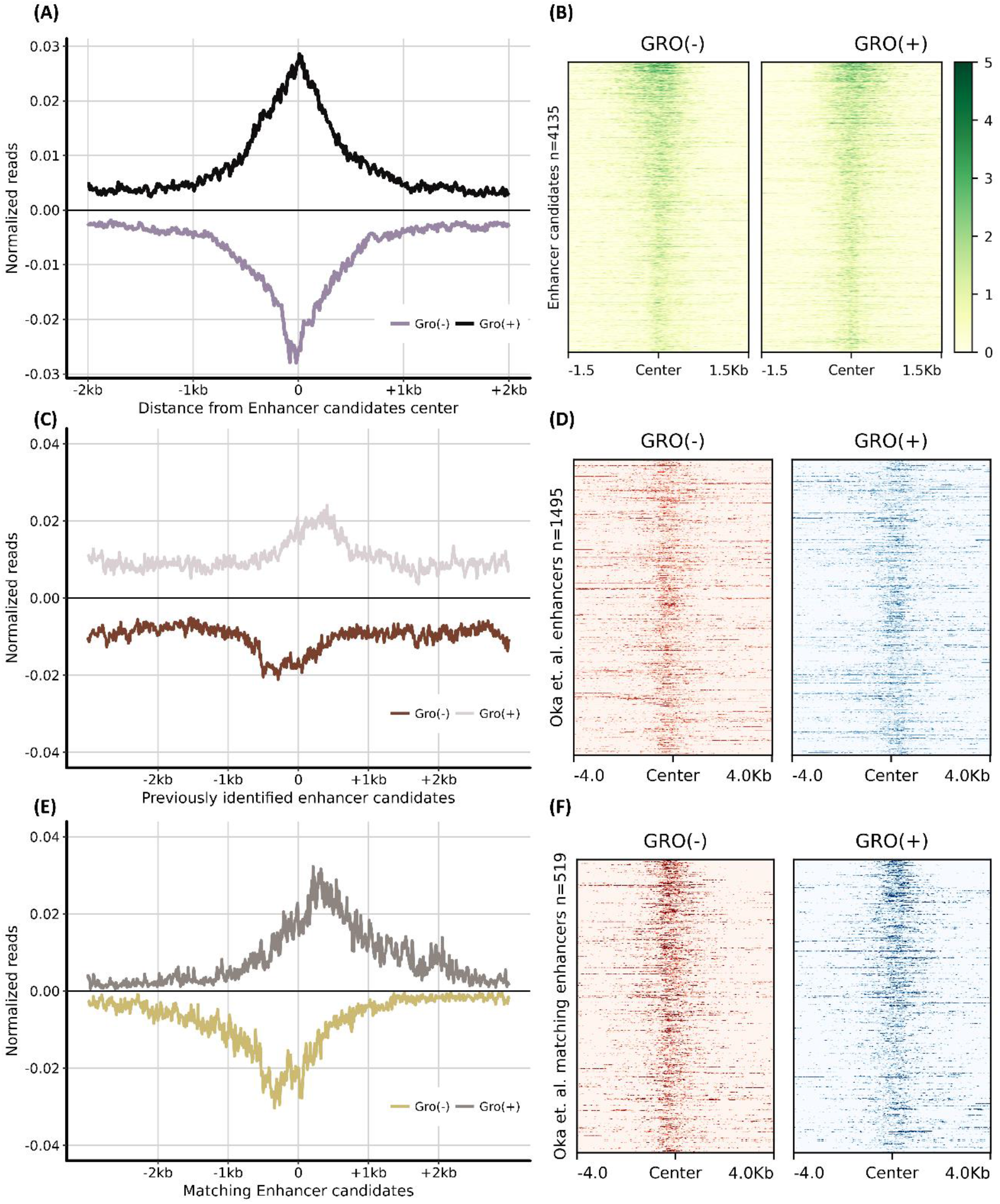
Enhancer candidates previously identified in Maize produce eRNAs. (**A**) GRO-seq reads^10^ mapping around maize enhancer candidates detected by dREG. Reads were sorted by strand and the normalize reads were plotted around the center of each candidate (n = 4135). (**B**) Heatmap representation of reads mapping to the enhancer candidate regions. The regions are sorted based on dREG scores. (**C**) GRO-seq reads were mapped to the regions previously identified as enhancer candidates by Oka et al.^9,30^ based on methylation, histone marks and chromatin accessibility (n = 1495). (**E**) A portion of the enhancer candidates (n = 519) reported by Oka et al. were also identified by dREG using just the GRO-seq reads. Heatmaps showing the GRO-seq signal of the regions in (C) and (E) are shown in (**D**) and (**F**) respectively.

Transcription of genomic enhancers was first described in 1992^31^, but the lack of adequate technology prevented further research on the subject until the late 2000’s^32,33^. While direct functions have been proposed for eRNAs as regulators of gene expression in metazoans^33,34^ there is no evidence of this in plants yet. Previous work in *Arabidopsis* did not identify eRNAs^9^, leading the authors to state that “if plants have enhancer elements, they rarely, if at all, produce transcripts.” Based on supporting research^12,35–37^, however, we believe the existence of plant enhancers is likely commonplace and independent of whether or not they are transcribed. Zhu et al.^37^ provided supporting evidence for this statement when they tested a small portion of nearly ten thousand enhancer candidates in *A. thaliana*: they validated 10 of the 14 enhancer candidates tested using a reporter assay. The GRO-arabidopsis data, however, did not show strong evidence of transcription in the regions identified by Zhu et al.^37^ (Figure S6). Additional data would be needed to fully validate the lack of enhancer transcription in *Arabidopsis*.

The results reported here suggest that plant transcriptional regulation may be more similar to that of mammals and other metazoans than previously thought. The lack of transcription previously observed in putative Arabidopsis enhancers may have been the result of relatively low read depths in the reported GRO-seq experiment. Further, the genome size of cassava is over six times that of *Arabidopsis*. Cassava therefore has much greater intergenic space, allowing for the identification of intergenic enhancer candidates whose expression is not confounded by that of nearby genes. The identification of intergenic transcribed enhancers in cassava and maize but not in *Arabidopsis* is consistent with the functional space hypothesis proposed by Mei et al.^38^ that simply predicts more functional genomic segments (e.g., enhancers) outside of genes in species with larger genomes.

## Acknowledgments

To the memory of Martha Hamblin who was a key part in the inception of this project. We would like to thank Deniz Akdemir from the Statistical Consulting Unit at Cornell, Karthik Ram for the Wes Anderson color palettes and Wojtek Pawlowski for help with the nuclei DAPI staining and visualization.

## Author contributions

R.L., D.P., B.Y., B.L, and G.B. performed experiments. R.L., D.P., J-L.J., J.L., and G.B. designed the experiment. E.B. provided intellectual input. R.L., G.B., J-L.J, J.L., D.P. contributed to writing the manuscript. All the authors reviewed the final manuscript.

## Data availability

The raw sequencing files have been submitted to the NCBI Gene Expression Omnibus (GEO) under accession GSE114758. All other data needed to evaluate the conclusions in the paper are present in the paper or the supplementary materials. Custom scripts for the NGS analysis, metagene plots and genomic partitioning, among others, have been made publicly available through the GitHub repository: https://github.com/tc-mustang/Pro-seq-Cassava.

## Materials and Methods

### Plant Materials and Nuclei Isolation

Cassava accession “Nase 3” (synonymous with “IITA-TMS-IBA30572” and “Migyera”) cuttings were grown in tubes containing enriched medium. Tubes were placed in growth chambers with 12 hours of light at 30°C for 6 weeks before tissue collection. Stem and leaves of approximately 25gr were ground with liquid nitrogen to a fine powder using a mortar and a pestle. The resulting powder was transfer to a coffee grinder containing cold 1X NIB buffer. Then, we used the CelLytica PN Plant nuclei isolation/extraction kit (Sigma-Aldrich) following the instructions for the “Highly-pure preparation of Nuclei.” The resulting solutions were frozen in liquid N2 and stored at -80° C. Most of the nuclei extraction protocol took place in a cold room (4° C) with all reagents on ice. A fraction of the nuclei preparations were stained with DAPI and visualized under a fluorescence microscope to test for concentration and nuclei integrity.

Maize inbred line B73 seeds were put into growth chambers. Shoots were collected 9 days after germination. Around 10 grams of plant tissue were ground with liquid nitrogen to a fine powder. Five grams of ground tissue was transferred into 50 ml fresh SEB extraction buffer (2.0% PVP, 10%TKE, 500mM sucrose, 4mM spermidine, 1mM spermine, 2.5% β-mercaptoethanol). Incubated on ice for 20 minutes, and then filtered through 2 layers of 100um nylon mesh. Triton X-100 was then added to a final concentration of 0.5% and incubated on ice for another 10 minutes. Then the lysate was centrifuged at 2000 rcf for 15 minutes at 4° C and the supernatant was recovered. The pellet was suspended in another 25 ml SEB extraction buffer and centrifuged again at 2000 rcf for 15 minutes. We added 10 ml nuclei storage buffer (50 mM Tris-Cl, 50% glycerol, 5 mM MgCl2, 0.1 mM EDTA, 0.5mM DTT) in the pellet and centrifuged at 2000 rcf for 5 minutes at 4° C. This step was repeated using 1 ml nuclei storage buffer. Finally, the pellet was resuspended and stored in 105 ul nuclei storage buffer. The protocol was conducted in a cold room (4° C).

### Pro-seq library preparation and sequencing

The PRO-seq protocol was performed as described by Mahat et al.^5^. Briefly, nuclei isolation washes away endogenous nucleotides, halting elongating RNA polymerases bound to chromatin. Precision run-on reactions were performed in the presence of equimolar amounts of unaltered ATP and GTP, as well as biotin-11-CTP and biotin-11-UTP (Perkin-Elmer). Notably, this two-biotin run-on will produce polymerase profiles with slightly reduced (~ 2-4 bp) resolution compared to a more typical four-biotin run on, given that RNA polymerases will primarily stall when incorporating the modified CTP and UTP^4^. RNA was extracted and base-hydrolyzed with NaOH. Hydrolyzed, biotin-labelled nascent RNAs were passed through a RNase-free P-30 spin column (Bio-Rad) and then enriched using M-280 streptavidin Dynabeads (Invitrogen). T4 RNA ligase 1 (NEB) was used to attach a 3’ RNA adaptor containing a six-nucleotide unique molecular index (UMI) for the removal of duplicate sequences produced by PCR (5’ - /5Phos/NNNNNNGAUCGUCGGACUGUAGAACUCUGAAC/Inverted dT/-3’). After a second biotin-enrichment, RNAs were submitted to RNA 5’ Pyrophosphohydrolase (RppH, NEB) treatment for 5’ decapping, and then 5’ phosphorylation with T4 polynucleotide kinase (T4 PNK, NEB), before ligation of the 5’ RNA adaptor (5’-CCUUGGCACCCGAGAAUUCCA-3’). Reverse transcription was performed with SSIII RT (Invitrogen) after a third biotin-enrichment. The cDNAs produced were PCR amplified for 13 cycles with Phusion polymerase (NEB) and size selected (120bp - 400bp) before sequencing. This protocol generated strand-specific libraries with every read starting from the 3’ end of the RNA. The RNA adapters used are TruSeq-compatible and libraries were reverse transcribed and amplified using primers from the Illumina TruSeq small RNA sequencing kit. Amplified libraries were assessed for quality on a bioanalyzer prior to sequencing on a HiSeq2500 with 100bp single reads.

### Analysis of NGS data

#### Processing and read alignment

The fastq files were scanned for any residual adapter sequence (5’ TGGAATTCTCGGGTGCCAAGG - 3’) using fastx_clipper from the FASTX_toolkit (http://hannonlab.cshl.edu/fastx_toolkit/), and the 3’ molecular barcode was removed. Reads were trimmed to a maximum length of 36bp, and the reverse complement was calculated because the Hiseq apparatus starts sequencing from the 5’ end. All downstream alignments were performed using Bowtie2^39^. The PRO-seq method described before is not exclusive to transcripts produced by the nuclear DNA Polymerase II, therefore the reads were aligned to the chloroplast genome to eliminate transcripts produced in this organelle. The remaining reads were mapped to the *Manihot esculenta* reference genome v6.1 (www.phytozome.com). Low-quality alignments were filtered and only reads mapping once to the genome were considered for further analysis (Table S1). Bedtools^40^ was used to get bedgraph files reporting only the number of 3’ end reads at each position. Finally, bigwig files were obtained from the bedgraph files using kentUtils (https://github.com/ENCODE-DCC/kentUtils).

#### Pro-seq read distribution

The percentage of the cassava genome transcribed was calculated using bedtools (Figure S7A). A saturation curve, which calculates the number of unique positions covered as a function of read depth was obtained using the bed-metric scripts (https://github.com/corcra/bed-metric.git) (Figure S7B). Normalized BigWig files representing the mapped reads were used to visualize each strand of the genome separately in the Integrative Genomics Viewer (IGV)^41^. The Metagene plots, histograms, peak scanning and gene expression values were generated using the HOMER software^42^ and meta-gene maker (https://github.com/bdo311/metagene-maker). Since the cassava genome is not readily available to work with HOMER, feature annotations were created separately, and the HOMER config files were modified. We approximated the TSS as the beginning of the 5’ UTR because the cassava genome annotation lacked precise TSS sites. The same approach was used for the Arabidopsis and Maize genomes.

#### Quantifying Pausing and Divergent Transcription

Genes were ranked based on their pausing index. The pausing indices were calculated, as previously described^34^. The average coverage in the promoter region (100 upstream of the Transcription Start Site (TSS) to 300 downstream of the TSS) divided by the average coverage of the gene body (300bp downstream of the TSS to the Polyadenylation site (PAS)). Divergent transcription indices were calculated similarly by taking the average coverage of the upstream promoter region (from 1kb upstream of the TSS to the TSS) in the antisense strand (with respect to the gene) divided by the average coverage of the TSS proximal region (300bp upstream the TSS to 300bp downstream the TSS) in the sense strand. We modeled the degree of promoter-proximal pausing and divergent transcription as measured by these indices by different factors including gene length, gene expression measured in RPKM (Reads per Kb of genic region per million mapped reads), cDNA length and number of exons using a linear model.

### Genomic Partitioning

Genomic Partitioning is a method to explore the genetic architecture of complex traits^21,22^. In this step, we calculated the heritability contribution from the enhancer candidate regions in the cassava genome and compared it with a random set of DNA regions of similar size and occupying a similar distribution across the cassava chromosomes.

#### Plant material and phenotypes

We analyzed data from the IITA cassava breeding program in Nigeria, including a fraction (689 clones, i.e., genetically unique individuals, each of which is clonally propagated) of the Genetic Gain (GG) collection, which comprises 709 elite and historically important clones. With these we analyzed 2302 clones developed as the cycle 1 of IITA’s Genomic Selection (GS) breeding program. In total, 3011 clones were used as source for phenotypes. For further details on the populations used, see Wolfe et al.^24,43^. We analyzed 5 traits: Dry matter content (DM), mean cassava mosaic disease severity (MCMDS), root number (RTNO), shoot weight (STWT) and fresh yield (FYLD). The phenotyping trials used in this study have been described before and all phenotype data is provided in the supplementary table S2.

#### Genotype Data

Genotyping-by-sequencing (GBS) libraries were prepared as described previously^44^. Marker genotypes were called using the TASSEL-GBS discovery pipeline^45^ using the *Manihot esculenta* genome assembly v6.1 (www.phytozome.net). The GBS markers were combined with the Cassava HapMap v2.0 variants from 241 deep sequenced cassava accessions^19^ to impute variants on all clones to whole genome sequence level in a single step with IMPUTE2^46,47^. The imputation procedure was performed as in Lozano et al.^48^ where the number of haplotypes used as the reference panel was set to 400, the effective population size (Ne) to 1000 and the imputation window to 5Mb. The resulting Oxford files where converted into the Plink^49^ binary format. In total, 3 million variants with a quality info score higher than 0.3 and Minor Allele Frequency (MAF) > 0.01 were obtained for the 3011 individuals used in this study.

#### Variance components estimation

Genomic partitioning analyses are imprecise in highly related populations because of high LD between partitions. We mitigated this problem by eliminating markers from the rest of the genome in high LD with the 9,665 cassava enhancer candidate regions (markers were removed that had allelic r^2^ > 0.9 and were closer than 100kb to enhancer markers). We also built 10 random sets of 9,665 regions with the same average length and approximate distribution in the genome as the enhancer candidates. As with the enhancer regions, random sets were forced to be outside any annotated gene (Figure S8), and markers from the rest of the genome in close physical distance and high LD were removed.

Genomic relationship matrices (GRMs) were calculated for focal (i.e., either enhancer or random sets) and ROG genomic partitions using the software LDAK5 following the ideas of Speed et al.^21^. Briefly, LDAK5 relationship matrices control short-range LD by assigning marker weights. Markers residing in low LD regions will have higher weights and are assumed to contribute more than markers in high LD regions. The LDAK5 model also assumes that a SNP’s heritability depends on its MAF, using an α value set to -0.25 as suggested in^21^. Finally, LDAK5 considers genotyping uncertainty as higher-quality observed markers should contribute more than poorly imputed markers. Genomic relationship matrices were calculated for the enhancer candidate partition and the rest of the genome partition. Separate analyses used Genomic relationship matrices based on the three random sets. Python scripts for these analyses can be accessed at the GitHub repository associated to this article.

The model fit to calculate the variance components was specified in matrix notation as:

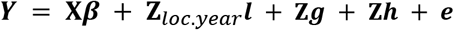

Where ***Y*** represents a vector with raw phenotypic observations, ***β*** represents the intercept and **X** is a vector of ones. The random effects include a single intercept for each location-year combination in which phenotypic trials took place and where ***ι*** ~ ***N***(0, ***Iσ***^*2*^_***ι***_) where ***I*** is the identity matrix and ***σ***^*2*^_***ι***_ is the associated variance component. The genetic variance components include ***g*** and ***h*** where ***g*** ~ ***N***(0, ***GRM_F_σ^2^_g_***) and ***h*** ~ ***N***(0, ***GRM_R_σ^2^_h_***). These two terms have known covariance structure calculated using LDAK5 for the focal (***GRM_F_***) or ROG (***GRM_R_***) partitions. The incidence matrices **Z**_***loc.year***_, and **Z** relate observations to the levels of trials and clones, respectively. Variance components were estimated using the ‘emmremlMultikernel’ function implemented in the ‘EMMREML’ R package^50^. There is no standard significance test for contrasting the alternative enhancer candidate hypothesis to the null model random set hypothesis (Deniz Akdemir, pers. comm.). Repeated sampling of the null model, however, shows non-overlap of its distribution of variance component estimates with the point estimate of the alternative for all four quantitative agronomic traits. We only sampled the null model ten times, creating ten random kernels because the two-GRM one-step model was computationally restrictively slow. Assuming independence among the four agronomic traits, the probability that all null models across all traits would explain less variance than the alternative, under the null hypothesis that random sets explain the same variance as enhancer candidates, would be (1 / 11)^4^ = 6.83e-5. In fact, root number and fresh root yield are strongly correlated (0.65 to 0.80,^51^) but are both uncorrelated to dry matter content and shoot weight. Thus, a conservative p-value for the hypothesis that enhancer candidates explain more variance in quantitative agronomic traits than random sets would be (1 / 11)^3^ = 7.51e-4.

## References

1. Adelman, K. & Lis, J. T. Promoter-proximal pausing of RNA polymerase II: emerging roles in metazoans. Nat. Rev. Genet. 13, 720–731 (2012).

2. Jennings, B. H. Pausing for thought: Disrupting the early transcription elongation checkpoint leads to developmental defects and tumourigenesis. Bioessays 35, 553–560 (2013).

3. Core, L. J., Waterfall, J. J. & Lis, J. T. Nascent RNA Sequencing Reveals Widespread Pausing and Divergent Initiation at Human Promoters. Science 322, 1845–1848 (2008).

4. Kwak, H., Fuda, N. J., Core, L. J. & Lis, J. T. Precise maps of RNA polymerase reveal how promoters direct initiation and pausing. Science 339, 950–953 (2013).

5. Mahat, D. B. et al. Base-pair-resolution genome-wide mapping of active RNA polymerases using precision nuclear run-on (PRO-seq). Nat. Protoc. 11, 1455–1476 (2016).

6. Sigova, A. A. et al. Divergent transcription of long noncoding RNA/mRNA gene pairs in embryonic stem cells. Proceedings of the National Academy of Sciences 110, 2876–2881 (2013).

7. Rennie, S. et al. Transcription start site analysis reveals widespread divergent transcription in D. melanogaster and core promoter-encoded enhancer activities. Nucleic Acids Res. (2018). doi:10.1093/nar/gky244

8. Core, L. J. et al. Analysis of nascent RNA identifies a unified architecture of initiation regions at mammalian promoters and enhancers. Nat. Genet. 46, 1311–1320 (2014).

9. Hetzel, J., Duttke, S. H., Benner, C. & Chory, J. Nascent RNA sequencing reveals distinct features in plant transcription. Proceedings of the National Academy of Sciences 113, 12316–12321 (2016).

10. Erhard, K. F., Talbot, J.-E. R. B., Deans, N. C., McClish, A. E. & Hollick, J. B. Nascent Transcription Affected by RNA Polymerase IV in Zea mays. Genetics 199, 1107–1125 (2015).

11. Law, J. A. & Jacobsen, S. E. Establishing, maintaining and modifying DNA methylation patterns in plants and animals. Nat. Rev. Genet. 11, 204–220 (2010).

12. Weber, B., Zicola, J., Oka, R. & Stam, M. Plant Enhancers: A Call for Discovery. Trends Plant Sci. 21, 974–987 (2016).

13. Kim, T.-K., Hemberg, M. & Gray, J. M. Enhancer RNAs: a class of long noncoding RNAs synthesized at enhancers. Cold Spring Harb. Perspect. Biol. 7, a018622 (2015).

14. Kim, T.-K. et al. Widespread transcription at neuronal activity-regulated enhancers. Nature 465, 182–187 (2010).

15. Danko, C. G. et al. Identification of active transcriptional regulatory elements from GRO-seq data. Nat. Methods 12, 433–438 (2015).

16. Wang, H. et al. CG gene body DNA methylation changes and evolution of duplicated genes in cassava. Proc. Natl. Acad. Sci. U. S. A. 112, 13729–13734 (2015).

17. Davydov, E. V. et al. Identifying a High Fraction of the Human Genome to be under Selective Constraint Using GERP++. PLoS Comput. Biol. 6, e1001025 (2010).

18. Cooper, G. M. et al. Distribution and intensity of constraint in mammalian genomic sequence. Genome Res. 15, 901–913 (2005).

19. Ramu, P. et al. Cassava haplotype map highlights fixation of deleterious mutations during clonal propagation. Nat. Genet. 49, 959–963 (2017).

20. Villar, D. et al. Enhancer evolution across 20 mammalian species. Cell 160, 554–566 (2015).

21. Speed, D. et al. Reevaluation of SNP heritability in complex human traits. Nat. Genet. 49, 986–992 (2017).

22. Yang, J. et al. Genome partitioning of genetic variation for complex traits using common SNPs. Nat. Genet. 43, 519–525 (2011).

23. Gusev, A. et al. Partitioning heritability of regulatory and cell-type-specific variants across 11 common diseases. Am. J. Hum. Genet. 95, 535–552 (2014).

24. Wolfe, M. D. et al. Prospects for Genomic Selection in Cassava Breeding. Plant Genome 0, 0 (2017).

25. Wolfe, M. D. et al. Genome-wide association and prediction reveals genetic architecture of cassava mosaic disease resistance and prospects for rapid genetic improvement. Plant Genome 9, (2016).

26. Jones, J. D. G. & Dangl, J. L. The plant immune system. Nature 444, 323–329 (2006).

27. Kremling, K. A. G. et al. Dysregulation of expression correlates with rare-allele burden and fitness loss in maize. Nature 555, 520–523 (2018).

28. Narita, T. et al. Human transcription elongation factor NELF: identification of novel subunits and reconstitution of the functionally active complex. Mol. Cell. Biol. 23, 1863–1873 (2003).

29. Booth, G. T., Parua, P. K., Sansó, M., Fisher, R. P. & Lis, J. T. Cdk9 regulates a promoter-proximal checkpoint to modulate RNA polymerase II elongation rate in fission yeast. Nat. Commun. 9, 543 (2018).

30. Oka, R. et al. Genome-wide mapping of transcriptional enhancer candidates using DNA and chromatin features in maize. Genome Biol. 18, 137 (2017).

31. Tuan, D., Kong, S. & Hu, K. Transcription of the hypersensitive site HS2 enhancer in erythroid cells. Proc. Natl. Acad. Sci. U. S. A. 89, 11219–11223 (1992).

32. Long, H. K., Prescott, S. L. & Wysocka, J. Ever-Changing Landscapes: Transcriptional Enhancers in Development and Evolution. Cell 167, 1170–1187 (2016).

33. Kim, T.-K. & Shiekhattar, R. Architectural and Functional Commonalities between Enhancers and Promoters. Cell 162, 948–959 (2015).

34. Chen, F. X. et al. PAF1, a Molecular Regulator of Promoter-Proximal Pausing by RNA Polymerase II. Cell 162, 1003–1015 (2015).

35. Shannon, S. et al. A Mutation in the Arabidopsis TFL1 Gene Affects Inflorescence Meristem Development. THE PLANT CELL ONLINE 3, 877–892 (1991).

36. Simpson, J. et al. Light-inducible and tissue-specific expression of a chimaeric gene under control of the 5’-flanking sequence of a pea chlorophyll a/b-binding protein gene. EMBO J. 4, 2723–2729 (1985).

37. Zhu, B., Zhang, W., Zhang, T., Liu, B. & Jiang, J. Genome-Wide Prediction and Validation of Intergenic Enhancers in Arabidopsis Using Open Chromatin Signatures. Plant Cell 27, 2415–2426 (2015).

38. Mei, W., Stetter, M. G., Gates, D. J., Stitzer, M. C. & Ross-Ibarra, J. Adaptation in plant genomes: Bigger is different. Am. J. Bot. 105, 16–19 (2018).

39. Langmead, B. & Salzberg, S. L. Fast gapped-read alignment with Bowtie 2. Nat. Methods 9, 357–359 (2012).

40. Quinlan, A. R., Quinlan & R., A. BEDTools: The Swiss-Army Tool for Genome Feature Analysis. in Current Protocols in Bioinformatics 11.12.1–11.12.34 (John Wiley & Sons, Inc., 2014).

41. Thorvaldsdottir, H., Robinson, J. T. & Mesirov, J. P. Integrative Genomics Viewer (IGV): high-performance genomics data visualization and exploration. Brief. Bioinform. 14, 178–192 (2013).

42. Heinz, S. et al. Simple Combinations of Lineage-Determining Transcription Factors Prime cis-Regulatory Elements Required for Macrophage and B Cell Identities. Mol. Cell 38, 576–589 (2010).

43. Wolfe, M. D., Kulakow, P., Rabbi, I. Y. & Jannink, J.-L. Marker-Based Estimates Reveal Significant Non-additive Effects in Clonally Propagated Cassava (Manihot esculenta): Implications for the Prediction of Total Genetic Value and the Selection of Varieties. G3 g3.116.033332 (2016).

44. Elshire, R. J. et al. A robust, simple genotyping-by-sequencing (GBS) approach for high diversity species. PLoS One 6, e19379 (2011).

45. Glaubitz, J. C. et al. TASSEL-GBS: a high capacity genotyping by sequencing analysis pipeline. PLoS One 9, e90346 (2014).

46. Howie, B., Marchini, J. & Stephens, M. Genotype Imputation with Thousands of Genomes. G3: Genes, Genomes, Genetics 1, (2011).

47. Howie, B. N., Donnelly, P., Marchini, J., Hardy, J. A. & Abecasis, G. A Flexible and Accurate Genotype Imputation Method for the Next Generation of Genome-Wide Association Studies. PLoS Genet. 5, e1000529 (2009).

48. Lozano, R. et al. Leveraging Transcriptomics Data for Genomic Prediction Models in Cassava. bioRxiv 208181 (2017).

49. Chang, C. C. et al. Second-generation PLINK: rising to the challenge of larger and richer datasets. Gigascience 4, 7 (2015).

50. Akdemir, D. & Okeke, U. G. EMMREML: Fitting Mixed Models with Known Covariance Structures. https://cran.r-project.org/package=EMMREML. R package version 3.1 (2015).

51. Okeke, U. G., Akdemir, D., Rabbi, I., Kulakow, P. & Jannink, J.-L. Accuracies of univariate and multivariate genomic prediction models in African cassava. Genet. Sel. Evol. 49, 88 (2017).

